# Cryo-EM of soft-landed *β*-galactosidase: Gas-phase and native structures are remarkably similar

**DOI:** 10.1101/2023.08.17.553673

**Authors:** Tim K. Esser, Jan Böhning, Alpcan Önür, Dinesh K. Chinthapalli, Lukas Eriksson, Marko Grabarics, Paul Fremdling, Albert Konijnenberg, Alexander Makarov, Aurelien Botman, Christine Peter, Justin L. P. Benesch, Carol V. Robinson, Joseph Gault, Lindsay Baker, Tanmay A. M. Bharat, Stephan Rauschenbach

**Affiliations:** Department of Chemistry, University of Oxford, Oxford OX1 3QU, UK; Kavli Institute for NanoScience Discovery, Dorothy Crowfoot Hodgkin Building University of Oxford, Oxford OX1 3QU, UK; Thermo Fisher Scientific, 1 Boundary Park, Hemel Hempstead, HP2 7GE, UK; Structural Studies Division, MRC Laboratory of Molecular Biology, Francis Crick Avenue, Cambridge, CB2 0QH, UK; Department of Chemistry, University of Konstanz, Konstanz, 78457, Germany; Thermo Fisher Scientific, De Schakel 2, 5651GH, Eindhoven, The Netherlands; Thermo Fisher Scientific, Bremen, 28199, Germany; Thermo Fisher Scientific, 5350 NE Dawson Creek Drive, Hillsboro, OR, 97124, USA; Department of Biochemistry, University of Oxford, Oxford OX1 3QU, UK

## Abstract

Native mass spectrometry (native MS) is a powerful technique that provides information on stoichiometry, interactions, homogeneity and shape of protein complexes. However, the extent of deviation between protein structures in the mass spectrometer and in solution remains a matter of debate. Here, we uncover the gas-phase structure of *β*-galactosidase using single particle electron cryomicroscopy (cryo-EM) down to 2.6 Å resolution, enabled by soft-landing of mass-selected protein complexes onto cold TEM grids and in-situ ice coating. We find that large parts of the secondary and tertiary structure are retained from solution, with dehydration-driven subunit reorientation leading to consistent compaction in the gas phase. Our work enables visualizing the structure of gas-phase protein complexes from numerous experimental scenarios at side-chain resolution and demonstrates the possibility of more controlled cryo-EM sample preparation.

**One Sentence Summary:** Electrospray ion-beam deposition on cold grids and in-vacuum ice growth enable cryo-EM of mass-selected proteins at 2.6 Å.

---

Over the past decade, native mass spectrometry (native MS) has evolved into a powerful tool for protein structural biology, because it provides information on homogeneity, stoichiometry, conformation, and interactions with ligands.^1–4^ Native MS is based on native electrospray ionization (ESI), in which intact proteins and protein complexes are transferred gently from solution into the gas phase by the application of low electrospray potential, using low flow rates, aqueous solutions, volatile buffers, and, where appropriate, addition of membrane mimetics.^5,6^ Since native MS aims to provide information relevant to proteins in their native, solvated environment, results are often interpreted in the context of X-ray crystallography or electron cryomicroscopy (cryo-EM) structures.^7–11^ However, since the early days of native MS, the extent to which the native protein fold and interactions are preserved upon transfer into the gas phase has been a matter of intense investigation and debate.^12–17^

Native MS comprises a range of analytical techniques that provide molecular structural information. Ion-mobility spectrometry can demonstrate retention of the globular protein shape, but also gas-phase collapse and unfolding can be observed, depending on charge state, degree of hydration, and temperature.^18–20^ Hydrogen-deuterium exchange reveals that similar amides are exposed or protected in solution and in the gas-phase.^21,22^ Various forms of ion activation show retention of native stoichiometry and conformation.^23^ In addition, molecular dynamics (MD) studies suggest that proteins adopt kinetically-trapped, compacted gas-phase structures that preserve the native secondary structure to a large extent.^17,24–29^ In particular, formation of additional intramolecular hydrogen bonds, also known as self-solvation, following dehydration has been proposed.^30^

Together, these findings suggest that near-native structures can be retained in the gas phase. However, due to the lack of spatial resolution in experimental MS data, the structural detail in MD simulations is difficult to validate at the atomic level and thus the degree to which biologically relevant structural motifs are retained in gas-phase protein ions remains unclear. Furthermore, this limitation has prevented access to a detailed molecular picture of the role of solvation in protein structure and folding.

To enable a detailed comparison of native solution structure and near-native gas-phase structure, soft-landing native electrospray ion-beam deposition (ESIBD) of protein ions directly onto cryo-EM grids has been suggested.^31^ In native ESIBD, ions produced by native ESI are mass-to-charge (*m/z*)-filtered and thermalized in a collision cell to obtain a molecular ion beam of narrow energy distribution (Fig. 1A). The ion-beam is then deposited with a very low deposition energy onto a substrate in vacuum. This approach has the potential to establish a direct link between chemical information from native MS and structural information from cryo-EM.

**Figure 1:**
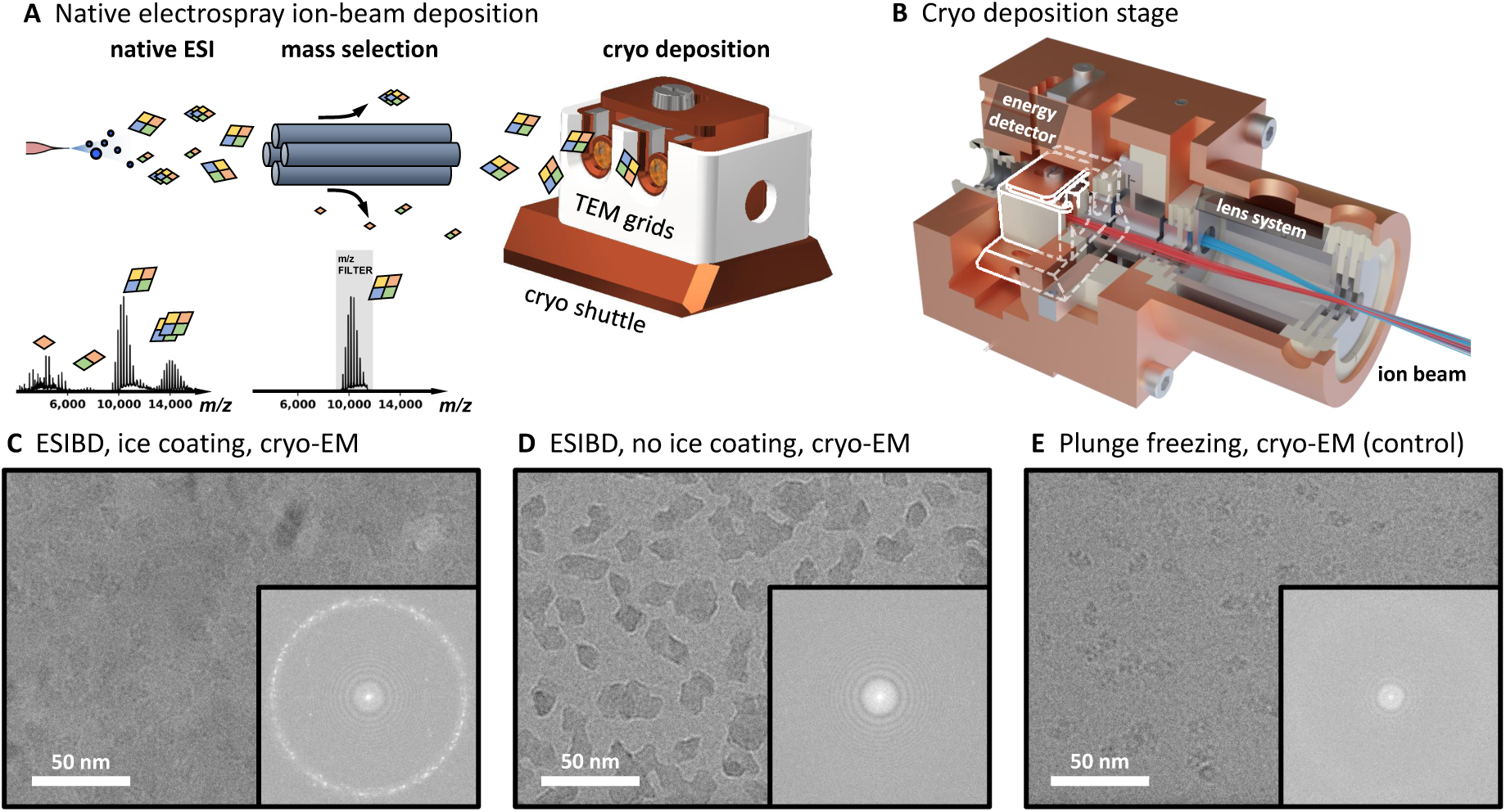
Determination of solution and gas-phase protein structure. **A** Na-tive electrospray ion-beam deposition workflow. Ions are transferred into the gas phase using native electrospray, mass selected, and deposited with controlled energy, density, and distribution on cold grids inside the cryo shuttle. Full and mass-selected spectra of *β*-galactosidase are shown. **B** Render of cryogenically-cooled landing stage. Protein ion beam is guided to the two grid positions using an electrostatic lens system. The stage is cooling grids and shielding them from contamination and radiation. The position of the cryo shuttle inside the cryo stage is highlighted in white. **C** Cryo-EM micro-graph from native ESIBD sample with ice-growth and corresponding power spectrum, showing non-vitreous ice due to temperature increase during transfer. **D** Cryo-EM micro-graph from native ESIBD sample without ice-growth and corresponding power spectrum, showing minimal ice. **E** Cryo-EM micrograph obtained from plunge-frozen sample and corresponding power spectrum showing vitreous ice.

Already, direct combination of native MS and various analysis methods revealed retention of globular shape, and biological activity of proteins and viruses.^32–35^ More recently, improved workflows using cryo-EM, negative stain transmission electron microscopy (TEM), or low-energy electron holography have demonstrated that protein complexes retain their overall shape at a resolution in the nanometer range.^35–40^ However, in the absence of a method that provides structures of proteins at the level of side-chain resolution, the extent to which the solution structure is retained in the native ESIBD process has remained unclear.

Here we present a 2.6 Å resolution cryo-EM density map of the tetrameric protein complex *β*-galactosidase, obtained from a sample prepared by in-vacuum deposition of a mass-selected, gas-phase protein ion beam onto a cryo-EM grid. This is enabled by our native ESIBD instrumentation, which delivers proteins at very low translational energy onto substrates held at cryogenic temperatures, coat them with ice grown from water vapor, and transfer them through a controlled inert atmosphere into liquid nitrogen. The three-dimensional (3D) map obtained from our data shows the structure of the adsorbed, gas-phase protein ion, which is remarkably similar to the structure obtained when using conventional, plunge-freezing sample preparation. The local fold and the integrity of the subunits are retained. Differences are observed at the protein surface and in the relative arrangement of the subunits. MD simulations suggest these changes correlate to compaction of previously water-filled cavities and grooves upon dehydration. Remarkably, the resolution obtained after averaging more than 400,000 single particle images of *β*- galactosidase suggest the formation of a single, well-defined, gas-phase structure under the conditions described herein.

## ESIBD workflow

We performed native ESIBD using, a previously described, modified, commercial mass spectrometer (Thermo Scientific^TM^ Q Exactive^TM^ UHMR instrument), which generates intense, *m/z* -filtered, and thermalized ion beams.^36,41^ This instrument was extended by a cryogenically-cooled deposition stage, shown in Fig. 1B (complete system shown in Fig. S1). TEM grids with 2 nm amorphous carbon films suspended on a holey film are held at cryogenic temperatures (T = 130 K) in ultra-high vacuum (p *<* 10*^−^*^9^ mbar). Compared to deposition at room temperature, cryogenic conditions suppress thermal diffusion and structural fluctuations immediately after deposition and during sample transfer.^42^

Up to two TEM grids were loaded and retrieved using a custom cryo shuttle, compatible with Thermo Scientific^TM^ Aquilos^TM^ hardware.^43^ Deposition began after optimizing *m/z* -selection, ion-beam intensity, and deposition energy.^36^ In contrast to the conventional plunge-freezing procedure, where grid, solvent, and proteins are frozen simultaneously, protein ions freeze individually as they land on the already cold substrate. After deposition, the shuttle passed from the vacuum of the deposition chamber through clean nitrogen gas into liquid nitrogen, without exposure to ambient conditions. During sample transfer, grids were shielded from heat and contaminants using a mechanical shutter mechanism within the shuttle (Fig. S2). To coat the cold samples with thin layers of ice, the partial pressure of water was temporarily increased while keeping the shuttle inside the stage.

## Structure determination

Using our native ESIBD workflow we prepared samples of *β*-galactosidase with and without coating in ice (see Fig. 1C and D). A third sample was prepared using the conventional plunge-freezing workflow, starting with the same protein solution (see Fig. 1E). The data was recorded on a Krios 300 kV cryo-TEM (Thermo Fisher Scientific) and processed using cryoSPARC^44^ (see SI).

All samples show similar particle shapes, density, and distribution. The ice-free native ESIBD sample provides significantly higher contrast than the those with ice. The absence of ice, and thus cleanliness of the sample transfer, is also confirmed by the absence of ice signals in the power spectrum. However, data from this sample did not result in high resolution, showing the molecular envelope with only rudimentary internal features and surrounded with a high-density shell (see Figs. S5, S6). Thus, we will focus on the comparison of the ice-coated, native ESIBD and plunge-frozen control samples in the following.

The corresponding micrographs in Figs. 1C and E show similar contrast. The presence of non-vitreous ice in the native ESIBD sample, revealed by diffraction spots in the power spectrum, indicates an increase of grid temperature to more than -140 C during sample transfer. Corresponding two-dimensional (2D) classes are shown in Fig. 2A and B. The 2D classes of the native ESIBD sample show significantly more detail compared to those obtained in previous studies where deposition and transfer were performed at room temperature.^37^ We find preferred orientation of particles in the native ESIBD sample, which likely reflects maximization of van der Waals interaction with the carbon substrate in vacuum. Clearly defined elements of the secondary structure are identified in the 2D classes of both samples. In particular the diamond shaped classes show features that directly correlate to the central *α*-helices and *β*-sheets at the tips. A reduction of overall size of the complex and the central cavity are the main differences observed in 2D between native ESIBD and plunge-frozen sample.

**Figure 2:**
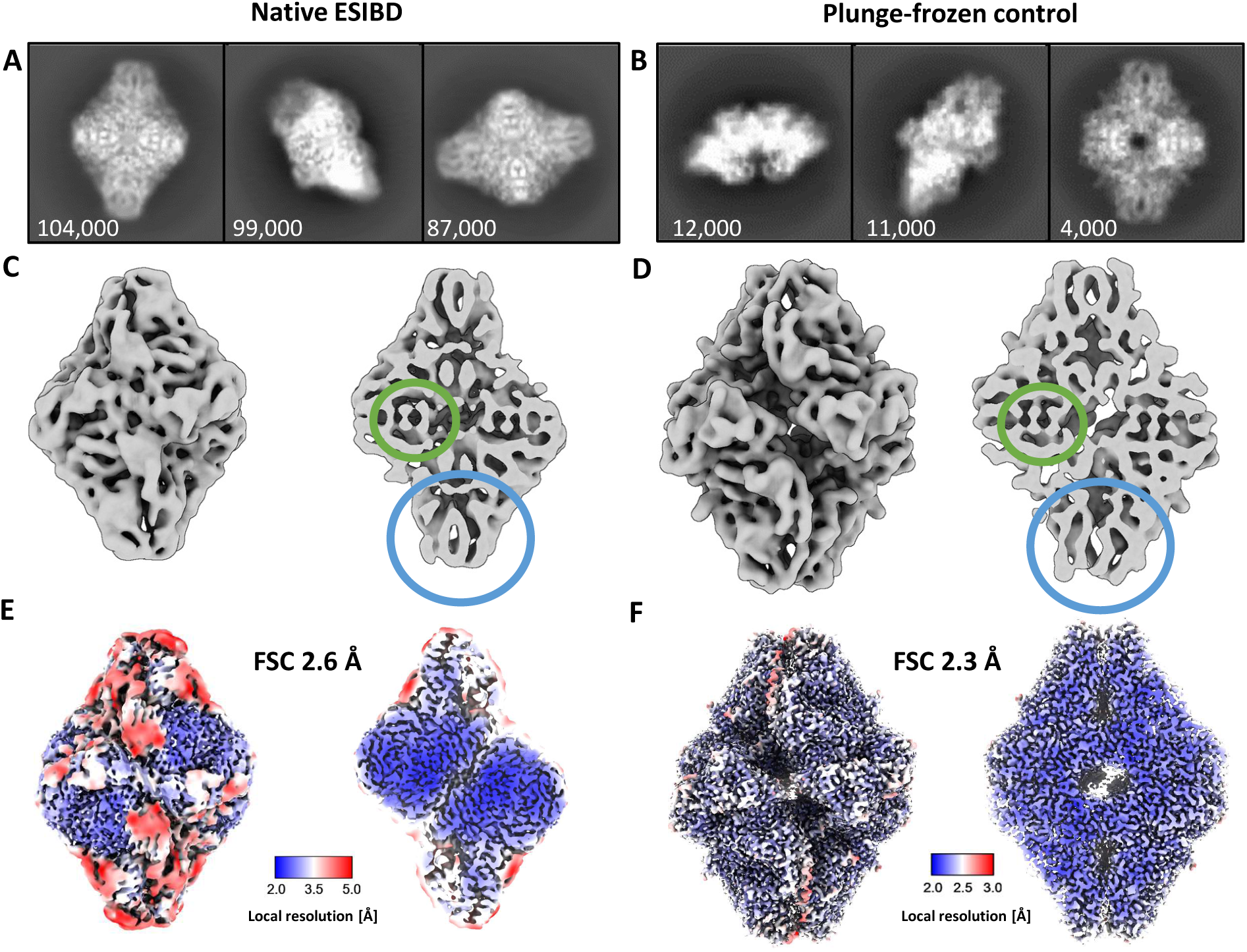
Comparison of 2D classes and 3D maps from native ESIBD and plunge-frozen control samples. **A**, **B** selected 2D classes (number of particles indi-cated). **C**, **D** Cryo-EM density maps (low-pass filtered to 8 Å to improve visibility of features) corresponding to the 2D classes above. *α*-Helices and *β*-sheets are highlighted by green and blue circles, respectively. The native ESIBD sample shows overall smaller size and reduced dimensions of grooves and the central cavity. **E**, **F** High-resolution den-sity maps (2.6 and 2.3 Å resolution estimated by gold-standard FSC). Color shows local resolution. Less resolved regions in the native ESIBD sample (panel **E**) coincide with typically solvated areas. Decreased resolution in these regions, though less pronounced, can also be seen for the plunge-frozen sample (panel **F**). Movie 1 shows an animation of the 2.6 Å map.

3D cryo-EM density maps (Fig. 2C-F) for the native ESIBD and the plunge-frozen control sample were obtained with a resolution of 2.6 and 2.3 Å, respectively. The 3D map for the control sample is in very good agreement with the previously published model PDB 6CVM.^45^ Both maps clearly show secondary structure, including *α*-helices and *β*- sheets (highlighted in Fig. 2C and D). Native ESIBD led to a compaction of about 10, 14, and 20 % along the long, short, and intermediate principal axis, respectively. The size of the central cavity is decreased by 40 %. Local resolution plots (Figs. 2E and F) indicate that the core regions of the subunits are highly ordered and resolved at below 3 Å, showing information on the side-chain level. However, surface-exposed areas that typically interact with solvent have poorer local resolution, suggesting local structural heterogeneity.

## Solution vs. gas-phase structures

Next, we investigated structural differences between the solution structure and the native ESIBD structure in more detail. By relaxing the *β*-galactosidase protein data bank (PDB) model 6CVM (Fig. 3C) into the map from native ESIBD using molecular dynamics flexible fitting in ISOLDE ^46^ we generated a tentative model, shown in Fig. 3A. It is more compact compared to the solution model, but the original secondary structure features are preserved and fit the core of the map well as seen by the color-coded cross-correlation. Movie 2 shows an animation of the original and relaxed model in the corresponding maps (low-pass filtered to 8 Å) as well as an extrapolated transition between them, visualizing the structural change during native ESIBD.

**Figure 3:**
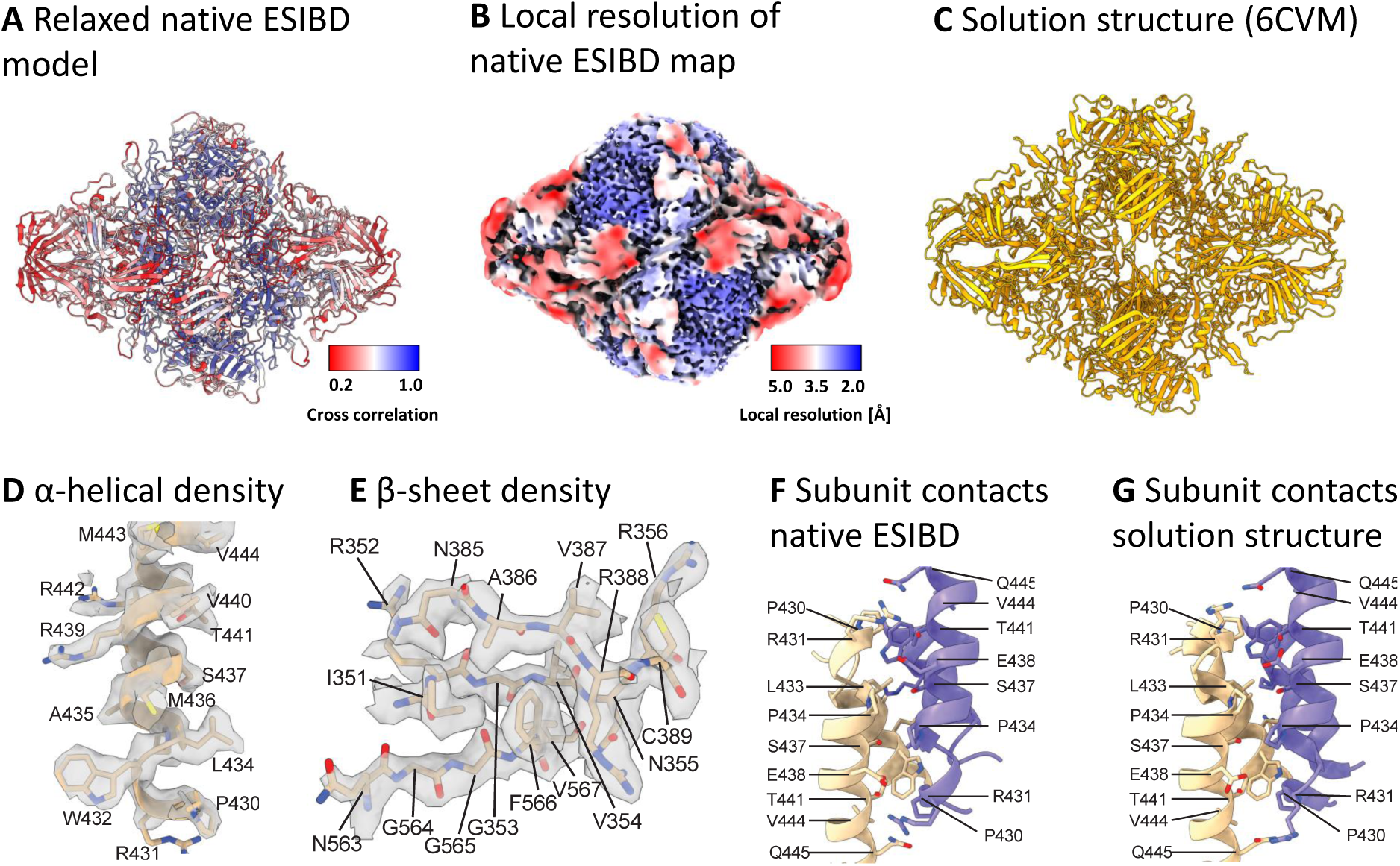
Atomic model of *β*-galactosidase from native ESIBD. **A** Relaxed full-length atomic model, created by molecular dynamics flexible fitting of the solution structure into the 2.6 Å native ESIBD map using ISOLDE. Color indicates goodness of fit of the relaxed model into the native ESIBD density, quantified by refinement of the atomic model against the density in Phenix and plotting the resulting per-residue cross-correlation value onto the model. Better fits for the model can be seen in the inner core region of the complex. **B** Color-coded local resolution map from native ESIBD, indicating strong correlation between resolution and quality of the relaxed model. **C** Solution structure (PDB 6CVM). **D,E** *α*-helical and *β*-sheet segment from a manually built atomic model within the density at 16 *σ* contour level. **F,G** Contacts between two helix pairs at the interface between two subunits in the built atomic versus the solution model. Side chains are shown for residues extending towards the interface.

While the surface-exposed areas of the map did not show sufficient side chain densities to enable model building, an unambiguous atomic model of the well-resolved core (300 residues) was built, with good agreement between map and model (model-vs-map FSC: 2.6 Å, see Table S1 and Fig. S7). The resulting model indicates retention of secondary and tertiary structure motifs. In particular, *α*-helices and *β*-sheets (Fig. 3D and E) and main chain segments (root mean square deviation (RMSD) below 1 Å; Fig. S8A and C) in the well-ordered monomer core show excellent local agreement between ESIBD and solution structure. Several contacts between subunits in the tetramer are retained from the solution structure (Fig. 3F and G). While one of the two subunit interfaces of each subunit indeed appears to be similar to the solution structure (RMSD 2.8 Å), the second one deviates more significantly (RMSD 6.6 Å), but consistently (see Fig. S8B). Taken together, this data suggests that the core of each subunit stays intact and highly ordered, while surface-exposed parts of the protein show reduced order. Overall, the architecture of the tetramer is maintained, but largely consistent variations in subunit interfaces cause a compaction of the complex, during which formerly water-filled cavities and grooves reduce in size.

## The effect of dehydration on structure

To investigate the effect of changing solvation during the ESI process, we performed MD simulations of the dehydration and rehydration of *β*-galactosidase (see SI for details). The simulations are based on the PDB model 6CVM,^45^ which represents the state of *β*-galactosidase captured in our map of the plunge-frozen control sample, and are independent from the experimental, native ESIBD cryo-EM density map. Fig. 4 shows the evolution of the radius of gyration (RoG) and number of intramolecular hydrogen bonds and compares models from simulation and experiment.

**Figure 4:**
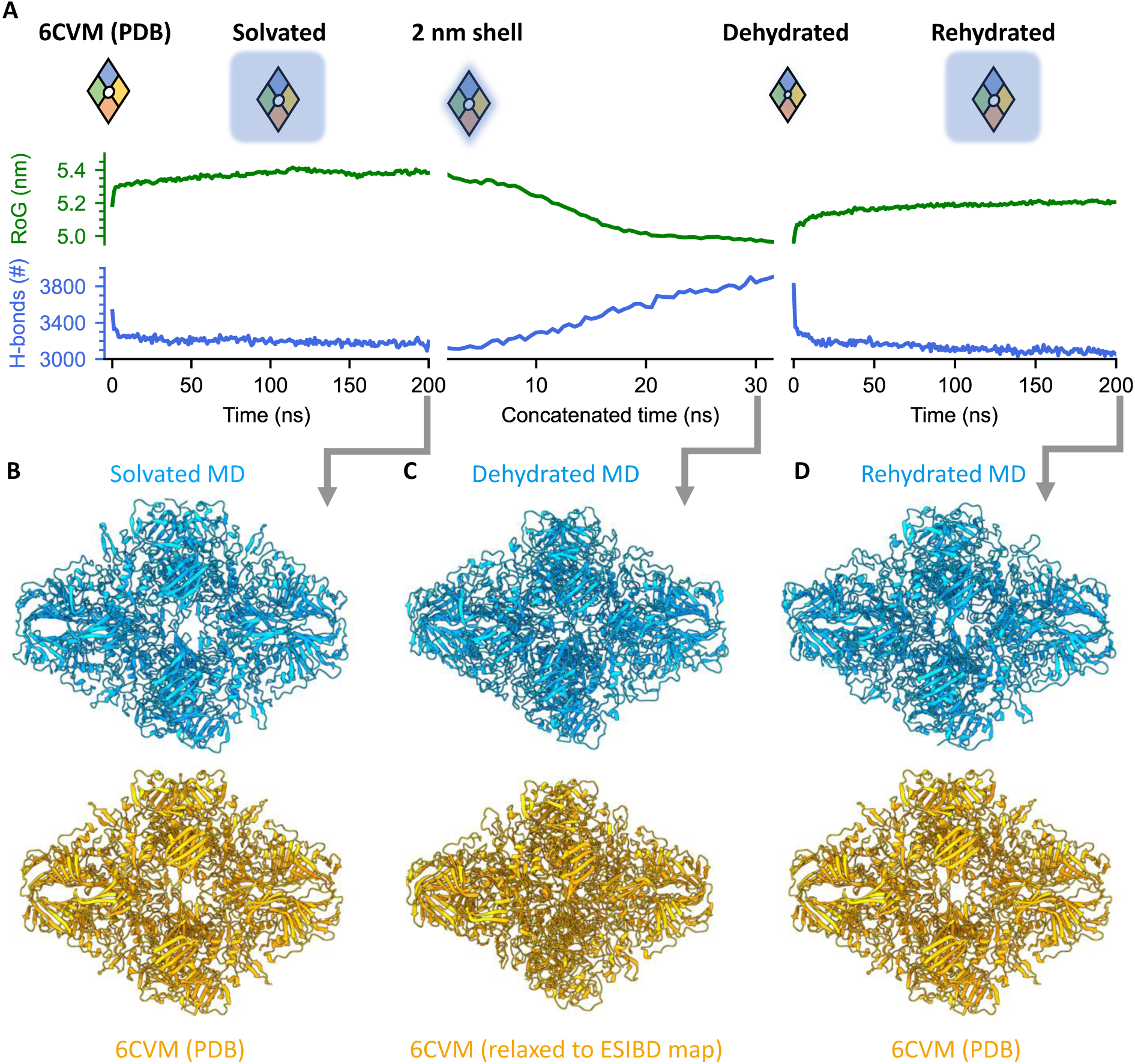
MD simulation of structural changes of *β*-galactosidase during solvation, dehydration, and rehydration. **A** Evolution of the radius of gyration (RoG, green) and number of intramolecular hydrogen bonds (blue) in the three phases of the MD simulation. Hydration state and size of *β*-galactosidase are indicated schematically. **B-D** Ribbon diagrams for models from simulation and experiment are shown in blue (top) and orange (bottom), respectively. **B** After solvation of 6CVM for 200 ns, only minor structural changes are observed. **C** The solvated model is gradually dehydrated, resulting in a compacted model, very similar to the native ESIBD model obtained from flexible fitting. **D** The dehydrated model was then allowed to rehydrate over 200 ns, largely recovering the secondary structure of 6CVM. See movie 3 for a morph and Table S2 for RMSD values between the models shown here.

First, we prepared a solvated initial model at 300 K (Fig. 4 left of A, B) after protonating 44 surface residues, corresponding to the dominating charge state observed in the experiment. Equilibration in solvent caused only minor changes to RoG, hydrogen bond number, and overall structure compared to 6CVM.

In order to generate a structural model of the gas-phase state of the protein, we dehydrated the initial structure at constant protonation state in short, free, atomistic simulations at 370 K following an adapted trajectory stitching approach (Fig. 4 middle of A, C).^47^ Here, in 64 simulation windows of 500 ps each, water molecules beyond a 2 nm shell around the protein surface were gradually removed until the amount of residual water converged to about 1400 molecules that could not be evaporated from the protein without increasing the simulation temperature. Importantly, at the end of the simulation there is no more hydration shell and the remaining water molecules are located inside the protein structure (see Fig. S9).

After dehydration, intriguingly, the outer dimensions and internal features - in particular grooves, central cavity, *α*-helices, and *β*-sheets - closely resemble the features of the native ESIBD cryo-EM density map. A decrease of the RoG by 7% and formation of 700 additional intramolecular hydrogen bonds indicate compaction and self-solvation, respectively.

Next, we simulated the rehydration of the dehydrated MD structure of *β*-galactosidase, motivated by previous work that has shown that rehydration of gas-phase structures, in silico or in experiments, can largely recover native structural motifs, collision cross sections, and biological activity.^14,24–27,41,48,49^ Rehydration over 200 ns (Fig. 4 right of A, D), led to restoration of the central cavity, outer dimensions, and recovery of many tertiary structure features of the native hydrated protein complex. The original number of intramolecular hydrogen bonds is recovered within the first 10 ns and keeps decreasing slightly afterwards, while the decrease in RoG is partially reversed. Movie 3 shows a morph between the three states shown in Fig. 4, highlighting the similarity of changes in overall size, extent of cavities and grooves, and subunit orientation.

## Discussion

Combining cryo-EM at side-chain resolution and molecular simulations allows us to understand how the structure of a protein is affected by the native ESIBD workflow. When water is removed, the protein compacts as a consequence of the relative motion of its subunits maximizing their mutual interaction. In this process, cavities and grooves that were previously filled with water are closed to form new intramolecular interfaces. Thus, structural changes are generally observed at the periphery of subunits where the molecule has been exposed to solvent before and is thus susceptible to self-solvation upon dehydration.^19,30,50,51^

Based on the close match of experimental and MD structures, we conclude that the observed change is mostly driven by dehydration, as it is the only effect considered in our MD simulations. Landing and substrate interaction are key parameters of the native ESIBD workflow and might contribute to the lower resolution at the protein surface. However, for *β*-galactosidase under native ESIBD conditions, i.e. 2 eV per charge deposition energy, on a thin carbon support, and cryogenic temperature, landing and substrate interaction contribute to a significantly smaller extent than dehydration to the overall structural changes.

Because it is based on averaging, single particle cryo-EM requires a chemically and structurally homogeneous set of single particle images to achieve high resolution. Our experiment shows that native ESIBD can produce samples with sufficient homogeneity to derive atomic models using cryo-EM at 2.6 Å resolution. The dehydrated gas-phase state is similar to the solution structure and in particular retains the local fold and secondary structure features. Our cryo-EM structural data represents a direct, experimental validation of structural, native mass spectrometry, which generally provides data at lower spatial resolution. Using rehydration in silico, we have shown that the native structure can largely be recovered from the dehydrated structure. Together with the partial atomic model, this demonstrates the possibility of determination of solvated protein structures from cryo-EM via native ESIBD, without the requirement of a priori knowledge beyond the protein sequence.

The high-resolution structural information obtained via native ESIBD will enable more detailed investigation of the effects of gas-phase activation, deposition energy, temperature, substrate and post-landing modifications steps. Further insights are to be expected from extending MD simulations to include landing and substrate interactions. Performance and information content of native ESIBD combined with cryo-EM may be improved by utilizing significantly higher ion transmission,^52^ ion mobility filtering,^53^ cross-linking,^54^ and low temperature membrane substrates tailored for efficient deposition energy dissipation, weak surface interaction, and low surface diffusion rates.^42,55,56^

In conventional cryo-EM sample preparation, poor sample quality is often directly linked to the plunge-freezing workflow. Thinning and freezing the solvent film can result in inconsistent ice thickness, ice-related radiation-induced motion, and denaturation at the substrate-solvent and air-solvent interfaces.^57–59^ Capitalizing on established MS workflows, native ESIBD has the potential to circumvent plunge-freezing related limitations of grid quality and preparation reproducibility, and allow for much higher control of sample purity, particle number, and particle distribution. Thus, native ESIBD may enable determination of structures of proteins that are not amenable to the conventional plunge-freezing workflow.

Comparison of our samples with and without ice underpins the importance of controlled modification of native ESIBD samples by the addition of water-ice or another matrix after landing. Compared to plunge freezing, the growth of ice layers can be controlled with much greater precision in vacuum to obtain optimal phase and thickness by regulating partial pressure of water vapor, growth temperature, and time.^60^ In particular, lower temperatures and increased transfer speed will allow to maintain vitreous ice, which has been previously demonstrated for the Aquilos sample transfer system.^43^ Other variations of the workflow, including matrix landing, protein hydration in the gas phase, and controlled rehydration after landing, may allow for a complete experimental route to native, hydrated protein structures, while retaining the advantages of gas-phase control.

## Acknowledgments

We acknowledge support from Thermo Fisher Scientific who provided the Q Exactive UHMR mass spectrometer within the framework of a technology alliance partnership and the Aquilos sample transfer system. We would also like to thank Sjors Scheres for critical comments. We thank Joseph Caesar, Rishi Matadeen, and Teige Matthews-Palmer for support with data collection and analysis.

## Author contributions statement

T.K.E., J.G., C.V.R., J.L.P.B., and S.R. conceived the experiments. P.F. and S.R. planned and implemented the custom deposition hardware. A.M. guided all hardware and software modifications of the Q Exactive UHMR mass spectrometer. A.K. suggested and coordinate the integration of the Aquilos sample transfer system. D.C. purified the protein sample. L.B. performed plunge freezing. T.K.E. and L.E. designed and implemented the cryo deposition stage. T.K.E. conducted solution preparation, native ESIBD, imaging, and data analysis for all other experiments with support from L.E., M.G., L.B., and J.B. J.B. and T.A.M.B. performed map validation, flexible fitting, and atomic model building. A.O and C.P performed MD simulations and analysis. T.K.E. and S.R. drafted the manuscript in consultation with J.B., T.A.M.B, A.O, C.P. All authors contributed to the interpretation of results and reviewed the manuscript.

## Data and materials availability

The 3D cryo-EM density map of the plunge-frozen control sample is available in the electron microscopy database (EMDB) under identifier EMD-18245. As it is equivalent to the map used for 6CVM, no additional model is provided. The 3D cryo-EM density map of the native ESIBD sample is available in the EMDB under the identifier EMD-18244. The atomic model of the core region is available in the PDB under identifier PDB ID 8Q7Y. All other data presented here does not comprise any new or high-resolution structures. Thus, they have not been uploaded to a repository. All generated and analyzed data sets are available from the corresponding author on reasonable request.

## 1 Supplementary Material

Materials and Methods

Supplementary Text

Figs. S1 to S9

Table S1 and S2

References^61–65^

Movie S1 to S3

## SUPPLEMENTARY INFORMATION

## 2 Materials and Methods

### 2.1 Protein preparation

*β*-galactosidase (G3153-5MG) was purchased from Sigma Aldrich. The lyophilized powder was resuspended in 200 mM ammonium acetate (pH 6.9) to a final concentration of 50 *µ*M.

For native ESIBD, it was desalted by passing through two P6 buffer exchange columns (7326222, Biorad), equilibrated with 200 mM ammonium acetate (pH 6.9) and diluted in 200 mM ammonium acetate (pH 6.9) to reach the concentration of 10 *µ*M and used without further purification.

For the preparation of plunge-frozen cryo-EM samples, it was desalted, purified, and transferred into a buffer (25 mM Tris, 50 mM NaCl, 10 mM EDTA, 2 mM MgCl_2_) using a Superdex 200 Increase 10/300 column (28990944, Cytiva life sciences).

Ammonium acetate (7.5 M, A2706-100ML) for native MS and buffer components for the plunge freezing solution, Tris (93362-250G), NaCl (S3014-500G), EDTA (DS-100G), MgCl2 (63068-250G), were also purchased from Sigma Aldrich. All concentrations are calculated with respect to the most abundant oligomers.

### 2.2 Native mass spectrometry

Borosilicate glass capillaries (30-0042, Harvard Bioscience) were pulled into nano electrospray ionization emitters, using a pipette puller (P-1000, Sutter Instrument), and gold coated using a sputter coater (108A/SE, Cressington). For native MS, up to 10 *µ*L of protein solution was loaded into an emitter and the emitter tip was clipped to produce an opening of 0.1 to 10 *µ*m.^62^ Electrospray ionization was initiated by applying a potential of 1.2 kV and a gentle nanoflow pressure (*<* 200 mbar above atm). Modifications of the ion source are described elsewhere.^36^

General instrument conditions were as follows: Source DC offset = 21 V, S-lens RF level = 200 (300 Vp-p), transfer capillary temperature = 60 °C, ion transmission settings set to ”High *m/z*” (700 Vp-p for the injection flatapole, and 900 Vp-p for the bent flatapole, transfer multipole, and collision cell), detector optimization ”High *m/z*”, injection flatapole = 5 V, interflatapole lens = 4 V, bent flatapole = 2 V, transfer multipole = 0 V, collision-cell pressure setting = 7 (N_2_), collision-cell multipole DC = -5 V, collision-cell exit-lens = -15 V.

For collection of mass spectra the instrument was operated in standard mode,^61^ and for ion deposition, a modified scan matrix was used that allowed ions to pass through the C-trap and collision cell directly into the deposition stage, without trapping. The number of water molecules attached to the protein complex ions under those conditions was estimated to be less than 200, by comparison to mass spectra after desolvation in the collision cell.^36^ The mass difference may also be explained by the presence of residual salt.

**Figure S1:**
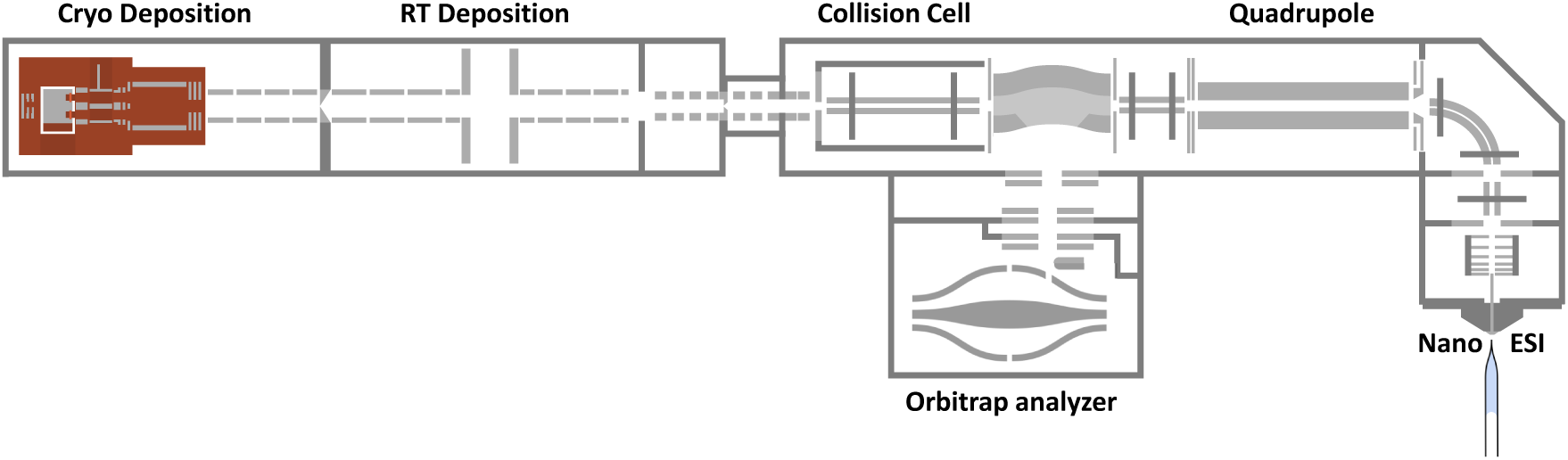
Overview of the electrospray ion-beam deposition (ESIBD) instrument. The ESIBD instrument consists of a commercial mass spectrometer (Thermo Scientific Q Exactive UHMR instrument, right) and a custom deposition setup (left). Proteins are transferred into the gas phase using a nano electrospray ionization source. They are then mass selected using a quadrupole mass filter and a mass spectrum is recorded using the Orbitrap^TM^ detector. For cryo-EM sample preparation, the ion beam is thermalized in the collision cell and guided using electrostatic ion optics though a differential pumping chamber into the cryo deposition stage (see Fig. S2A).

### 2.3 Preparation of the control grid

The control sample was prepared using a grid with mesh size 200, R2/1 holey carbon without amorphous carbon from Quantifoil. 3 *µ*L of a 2.5 *µ*M solution was applied to the grid, followed by blotting and plunging into liquid ethane, using a manual plunger.

### 2.4 Preparation of the native ESIBD grid

Grids were prepared and transferred as described in the main text. Fig. S1 shows an overview of the deposition instrument described in more detail elsewhere.^36,41^ Fig. S2 shows the cryo deposition stage and Aquilos transfer system including transfer rod, load lock, and preparation pot. At the beginning of a deposition experiment, clipped TEM grids are loaded into the shuttle at room temperature. Here, copper TEM grids from Quantifoil (C2-C15nCu40-01) with mesh size 400, R2/1 holey carbon covered with 2 nm amorphous carbon were used. The shuttle is loaded into the stage a few minutes before deposition, to allow for thermalization to the stage, which was held at 130 K using a liquid nitrogen heat exchanger. While inserting the shuttle, a spring-loaded mechanism exposes the grids for deposition and makes electrical contact to control the electric potential of and monitor ion current on the grids. The stage geometry prevents a direct line of sight between grids and warm surfaces to minimize contamination and radiative heating. The transfer rod is retracted to prevent heat transfer to the shuttle and to close the load-lock valve to obtain a cleaner vacuum. The pressure in the deposition chamber is 10*^−^*^9^ mbar, however, the pressure within the stage will be substantially lower, as it acts as a cryo pump.

The beam energy is measured using a retarding-grid energy detector. The deposited charge is monitored by a picoampmeter to control protein coverage. Protein ions are deposited with a deposition energy of 2 eV per charge, by applying a corresponding potential to the TEM grids. Close to monolayer coverage in the grid center is typically achieved in 30 minutes, and lower densities for efficient data collection can be found further from the center if needed. The second sample can be prepared using the same conditions or after varying the analyte, *m/z* -window, or deposition energy. After completion, transfer into liquid nitrogen is done in less than two minutes as described in Fig. S2.

**Figure S2:**
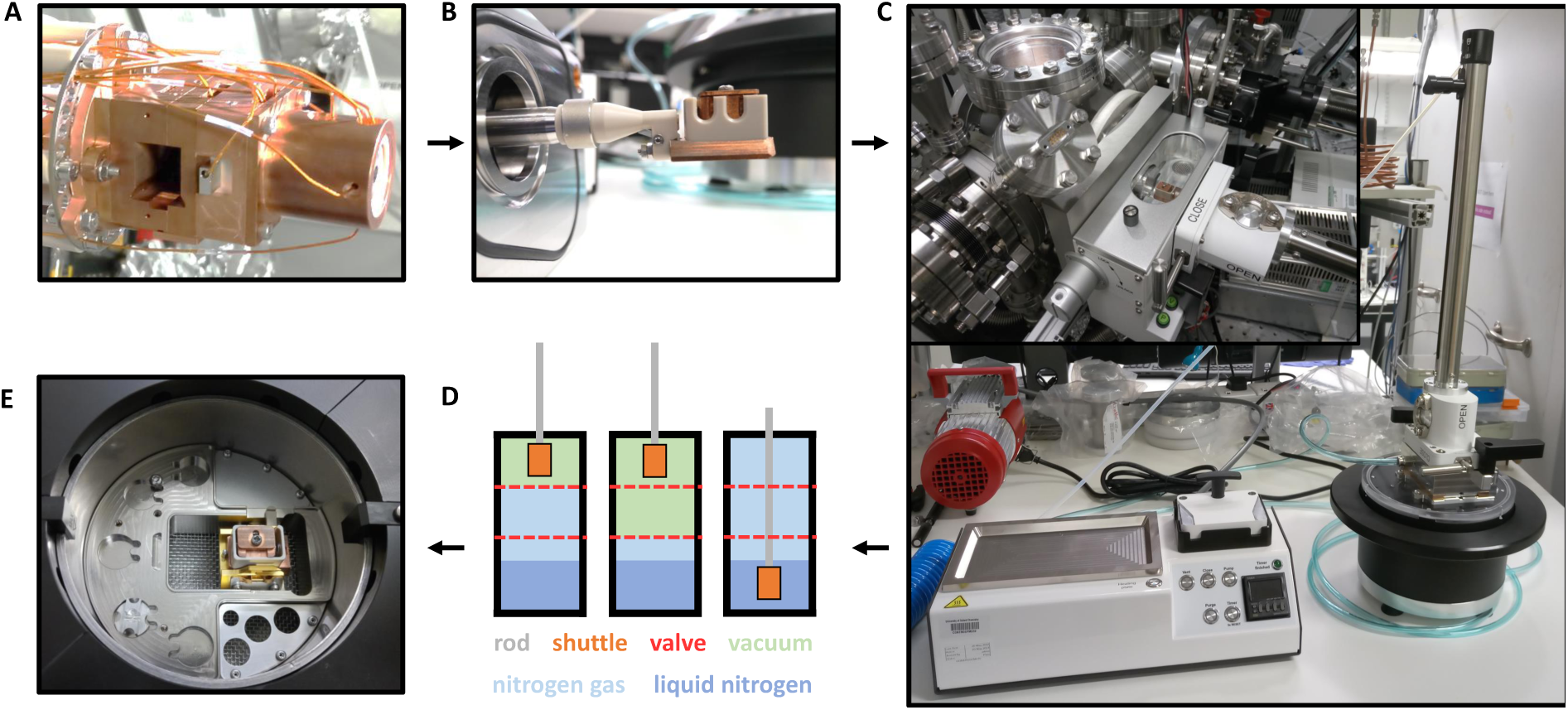
Overview of cold and clean transfer using Aquilos components. **A** Picture of the cryo deposition stage. **B** Picture of the cryo shuttle. Grids are shielded by the PEEK housing during transfer. **C** Picture of sample transfer rod attached to preparation pot. The lower left shows the preparation station, used to control pumping, venting, and keeping grid handling equipment dry. The insert shows the load lock attached to the deposition chamber. **D** Schematic of cold sample transfer. Left: Cold shuttle in vacuum of transfer rod. Preparation pot and load lock are purged with clean nitrogen gas. Middle: Evacuating load lock. Right: Load lock and transfer rod are vented with clean nitrogen gas. Shuttle is plunged into liquid nitrogen. **E** Picture of shuttle inside preparation pot. Grids are transferred from shuttle to conventional storage boxes under liquid nitrogen (not shown.)

### 2.5 Movie acquisition and processing

All micrographs were collected using a Thermo Scientific^TM^ Krios 300 kV cryo-TEM equipped with a BioQuantum^QR^ energy filter operated at a slit width of 20 eV and a K3 direct electron detector (both Gatan), located at the COSMIC Cryo-EM facility at South Parks Road, University of Oxford, UK. Automated data acquisition were controlled using EPU^TM^ software (Thermo Fisher Scientific). All movies were recorded in the tif format, using a range of defocus settings between -1 and -2.5 *µ*m, an exposure of 34 *e*^−^/Å^2^, and a magnification of 105,000 corresponding to a pixel size of 0.83 Å.

Data was processed using cryoSPARC.^44^ After running Patch Motion Corr. and Patch CTF jobs, particles were picked using template picking and extracted in 416*×*416 pixel^2^ boxes scaled down by a factor of 2. After multiple rounds of 2D and 3D classification, particles were re-extracted without down scaling and final maps were produced using Non-Uniform Refinement using Ab-Initio initial volumes. This was followed by Local Resolution Estimation and Local Filtering. D2 symmetry was imposed in all steps, but using C1 symmetry resulted in only slightly lower resolution. Figures and movies of the resulting 3D EM density maps were generated using ChimeraX.^63^ Further movie acquisition and processing details for the two samples compared in the main text is specified in Fig. S3, Fig. S4, and table S1.

**Movie 1**: Animation of the 2.6 Å native ESIBD map.

**Movie 2**: Animation of maps from native ESIBD (2.6 Å) and plunge-frozen control (2.3 Å), corresponding models and morph between models.

**Movie 3**: Morph between initial, dehydrated, and rehydrated MD models compared to corresponding experiment based solvated and relaxed PDB models.

**Figure S3:**
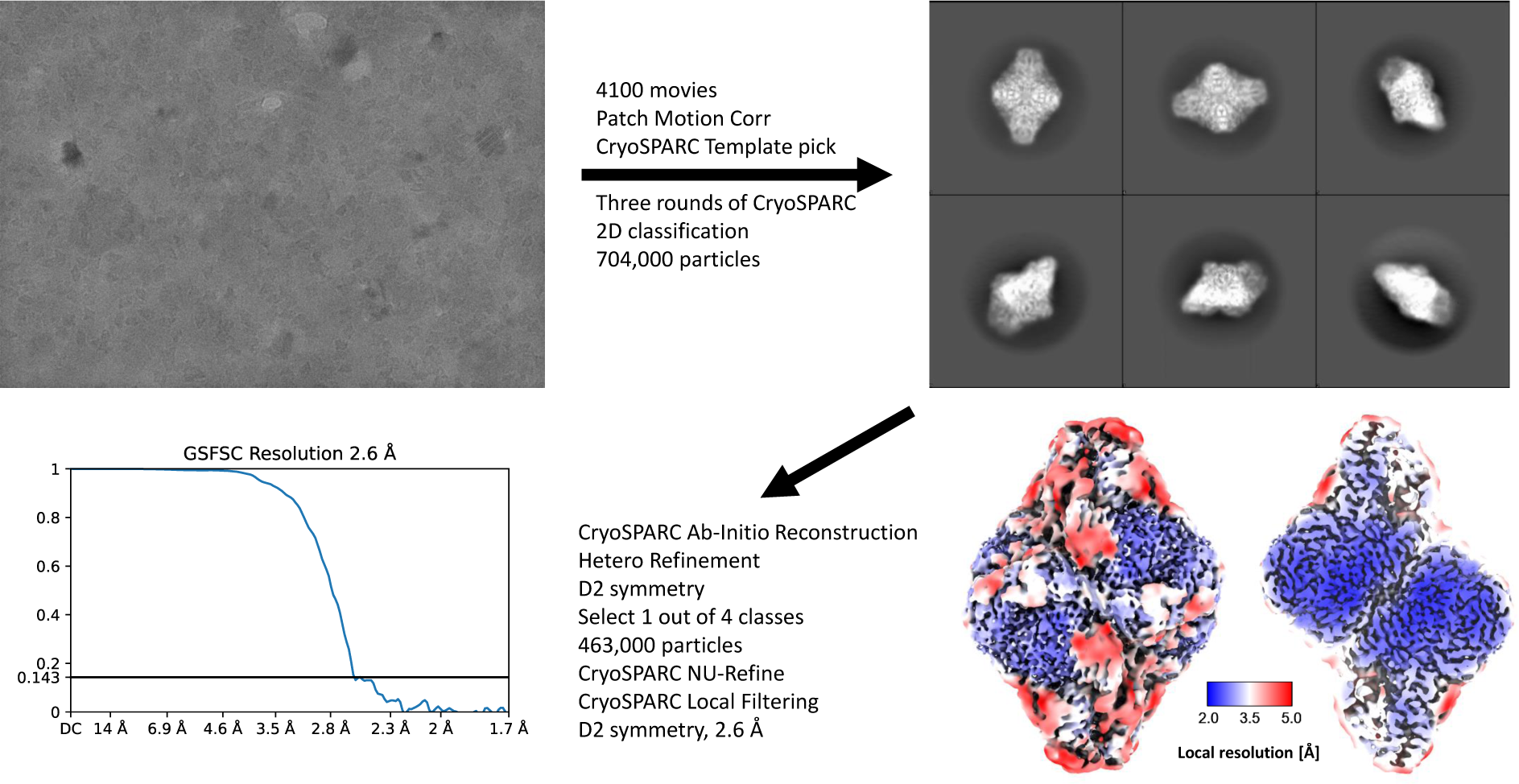
Data processing schematic for *β*-galactosidase from ice-coated native ESIBD sample. Using homogeneous refinement in CryoSPARC resulted in a 3.4 Å map. By using non-uniform refinement in CryoSPARC the resolution could be further improved to 2.6 Å.

**Figure S4:**
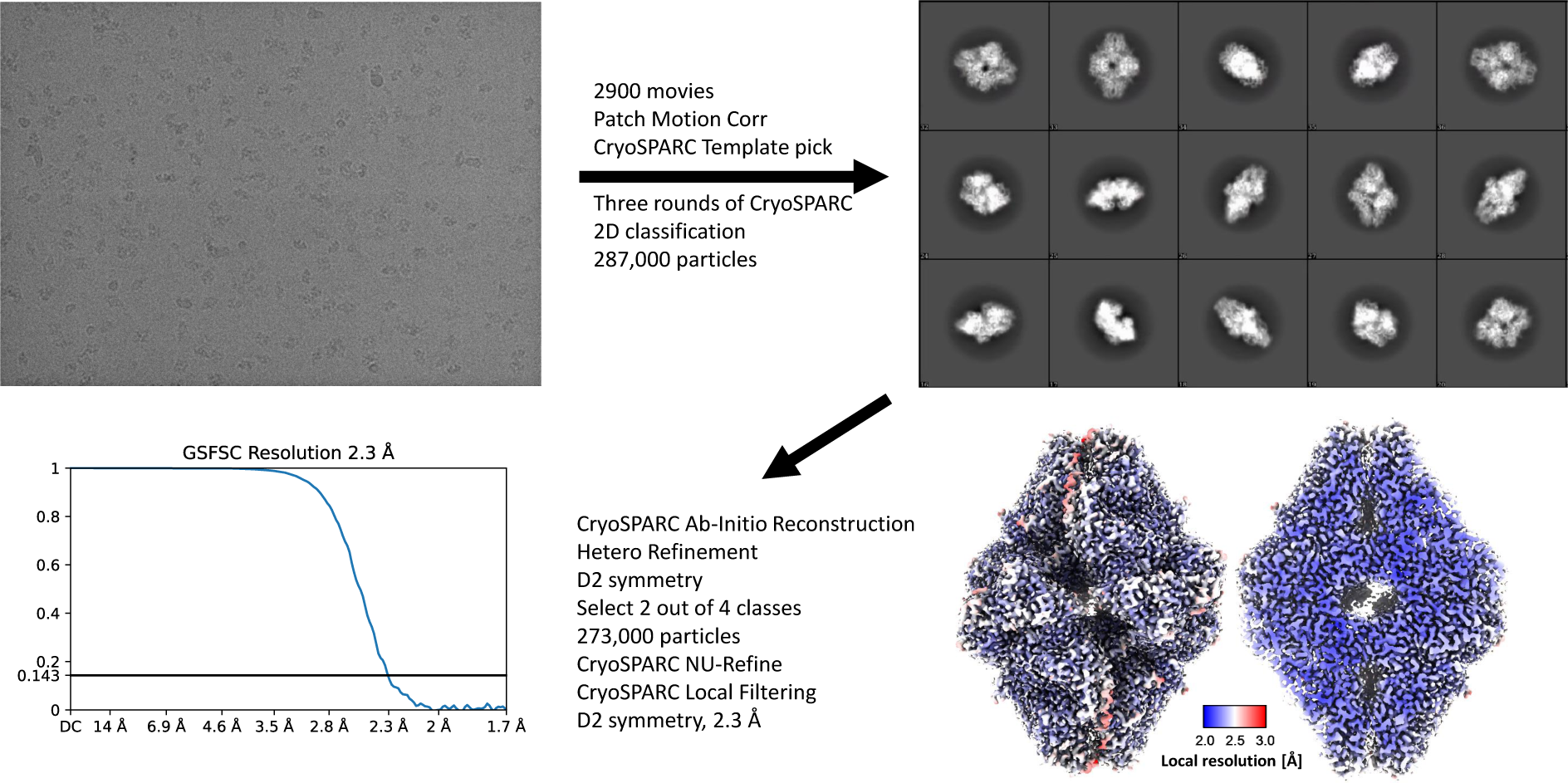
Data processing schematic for *β*-galactosidase from plunge-frozen control sample.

**Table S1:** *β*-galactosidase cryo-EM data acquisition and processing statistics.

| Data collection and processing | β-galactosidase native ESIBD | β-galactosidase solution structure |
| --- | --- | --- |
| Microscope | Krios G3 | Krios G3 |
| Magnification | 105,000 | 105,000 |
| Voltage (kV) | 300 | 300 |
| Electron exposure ($e^-/\text{\AA}^2$) | 34 | 34 |
| Defocus range ($\mu\text{m}$) | -1 to -2.5 | -1 to -2.5 |
| Pixel size ($\text{\AA}$) | 0.83 | 0.83 |
| Symmetry imposed | D2 | D2 |
| Initial particle images (no.) | 704,000 | 287,000 |
| Final particle images (no.) | 463,000 | 273,000 |
| Map resolution ($\text{\AA}$) | 2.6 | 2.3 |
| FSC threshold | 0.143 | 0.143 |
| Map resolution range ($\text{\AA}$) | 2.4 to 5.0 | 2.1 to 3.0 |
| Refinement |  | No atomic model built |
| Initial model used (PDB code) | 6CVM |  |
| Model resolution ($\text{\AA}$) | 2.2/2.5/2.6 | |
| FSC threshold | 0/0.143/0.5 |  |
| Map sharpening B factor ($\text{\AA}^2$) | -100 | -84 |
| Model composition |  |  |
| Non-hydrogen atoms | 10048 |  |
| Protein residues | 1212 |  |
| Ligands | 8 Mg, 8 Na |  |
| B factors ($\text{\AA}^2$) | | |
| Protein | 66.88 |  |
| Ligand | n/a |  |
| R.m.s. deviations |  |  |
| Bond lengths ($\text{\AA}$) | 0.002 | |
| Bond angles ($^\circ$) | 0.547 | |
| Validation |  |  |
| MolProbity score | 2.43 |  |
| Clashscore | 13.05 |  |
| Poor rotamers (%) | 3.77 |  |
| Ramachandran plot |  |  |
| Favored (%) | 94.62 |  |
| Allowed (%) | 5.38 |  |
| Disallowed (%) | 0 |  |

### 2.6 Ice-coated vs. ice-free native ESIBD samples

A third map was obtained from an ice-free native ESIBD sample as shown in Fig. S5. The quality of this map is significantly lower than the one obtained from the ice-coated sample. A high density at the edges of the map conceals the underlying protein structure. The outer surface of this high-density region is significantly larger than the map obtained from the ice-coated native ESIBD sample. It may be caused by the presence of an irregular ice shell of 2 to 4 monolayers around particles. Given the pressure (10*^−^*^7^ mbar), temperature (-140 °C), and time spend in the vacuum chamber (up to 4 hours), growth of a few monolayers of ice is very plausible. Alternatively, a data processing artifact caused by the high contrast of ice-free samples could be the cause. In both cases, adaptation of data analysis software might allow to obtain more information from such samples.

**Figure S5:**
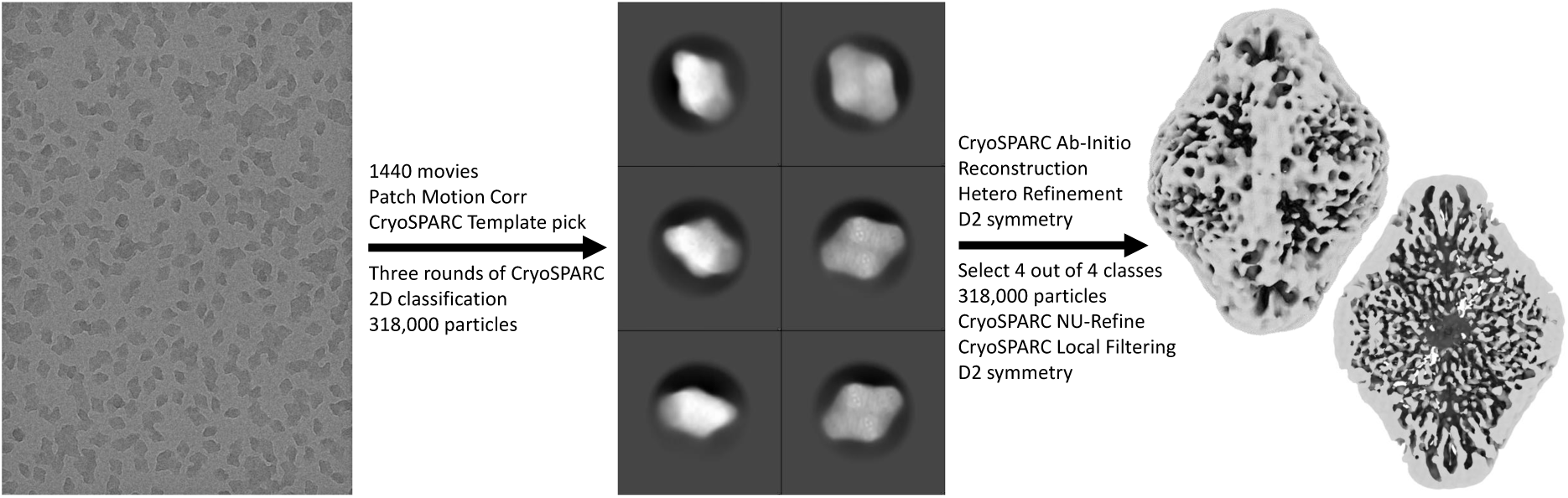
Data processing schematic for *β*-galactosidase from ice-free native ESIBD sample. A high density at the edges of the map conceals the underlying protein structure. The map only shows the molecular envelope with rudimentary internal features.

Better control of the amount of ice on native ESIBD samples will help to understand this effect. However, we have shown that it can be avoided in practice by ice-coating, which provides a homogeneous background similar to conventional plunge freezing samples. It is important to note that this step does not constitute rehydration, but only coating with additional ice grown from water vapor. As opposed to ice formation during plunge freezing, this does not expose particles to an air-solvent interface and particles are retained in the orientation and hydration state they were frozen in. Adding ice gradually from individual gas-phase water molecules provides the option to control the thickness of the ice layer and prevent buildup of tension in the ice.

Fig. S6 shows a direct comparison of the three maps discussed so far, and a map generated from our previous work, but with processing in cryoSPARC analogous to the other shown maps.^36^ The latter was made using a sample deposited at room temperature, followed by manual transfer through ambient conditions and plunging into liquid nitrogen. The first row shows 8 Å low-pass filtered maps while the second row shows corresponding full resolution maps. While the 8 Å maps of the control and ice-coated ES-IBD sample differ in overall size and size of cavities, they show similar secondary structure features as discussed in the main text. In contrast, the 8 Å maps of both ice-free samples only show rudimentary secondary structure features and exhibit the high-density shell discussed above, despite the differences in sample preparation workflow. Looking at the corresponding full resolution maps, the map from the sample prepared by landing on a cryogenically cooled grid contains more detail, though it is not sufficiently resolved to be used for model building. While landing on cryogenically cooled surfaces allowed us to obtain the 2.6 Å native ESIBD map, these observations emphasize that temperature is only one of many parameters that need to be carefully controlled to further improve the quality of native ESIBD samples.

**Figure S6:**
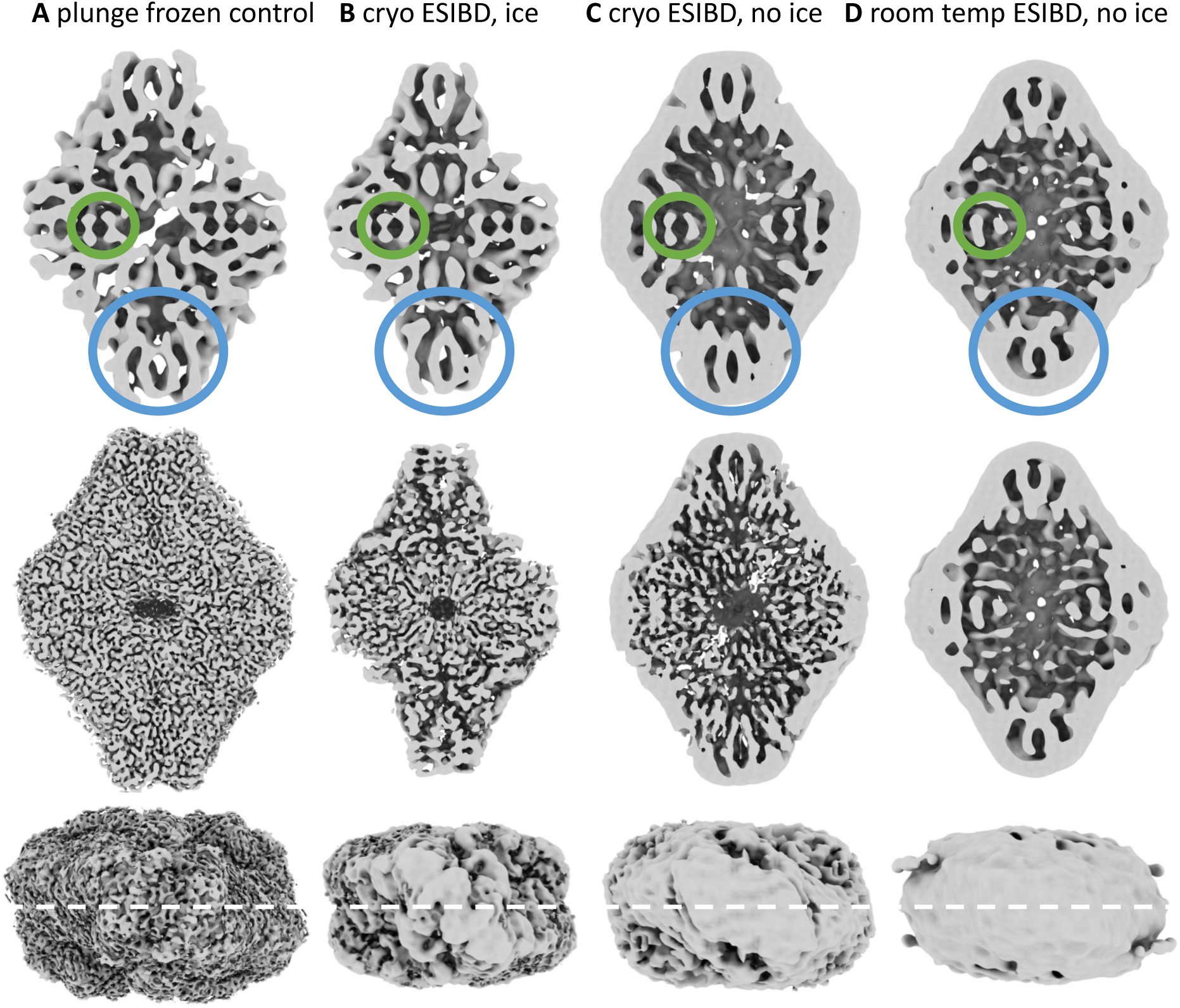
Comparison of maps for control, ice-free, ice-coated, and ice-free room-temperature deposition samples. The first row shows 8 Å low-pass filtered and the second row full resolution maps. The third row shows corresponding top views with a white dashed line indicating the section plane used for the sections above. The location of four characteristic central *α*-helices and four peripheral *β*-sheets are marked with green and blue circles in all 8 Å maps. Maps in panels **A** and **B** are control and ice-coated native ESIBD sample, already shown in Fig. 2 of the main text. **C** Map of the ice-free native ESIBD sample. **D** Map of an ice-free native ESIBD sample where deposition was done at room temperature.^36^

### 2.7 Model building

A previously published model of *β*-galactosidase (PDB 6DRV) was used as a starting model. Fitting into the cryo-EM map obtained by conventional plunge-freezing reveals good agreement with this model. Fitting into the ESIBD cryo-EM map, however, revealed significant differences between this model and the ESIBD cryo-EM map. In order to obtain a starting model for model building, non-ionic ligands and water molecules within the 6CVM model were removed, and the model was relaxed into the ESIBD map using ISOLDE, resulting in good agreement of secondary structure elements.

In order to obtain an atomic model of the resolved parts of beta-galactosidase, the relaxed model was first manually inspected and trimmed by removing parts that had poor agreement with the density and/or consistently showed no obvious side chain densities. The resulting trimmed model was inspected in Coot, where segments that did not fit their density were deleted and manually rebuilt using large aliphatic side chains as anchors. The model was then subjected to multiple rounds of real space refinement in PHENIX and manual building. Some residues, often near disordered regions, had poor side chain densities and resulted in rotamer outliers. The statistics and map presented are the result of real space refinement in PHENIX.

**Figure S7:**
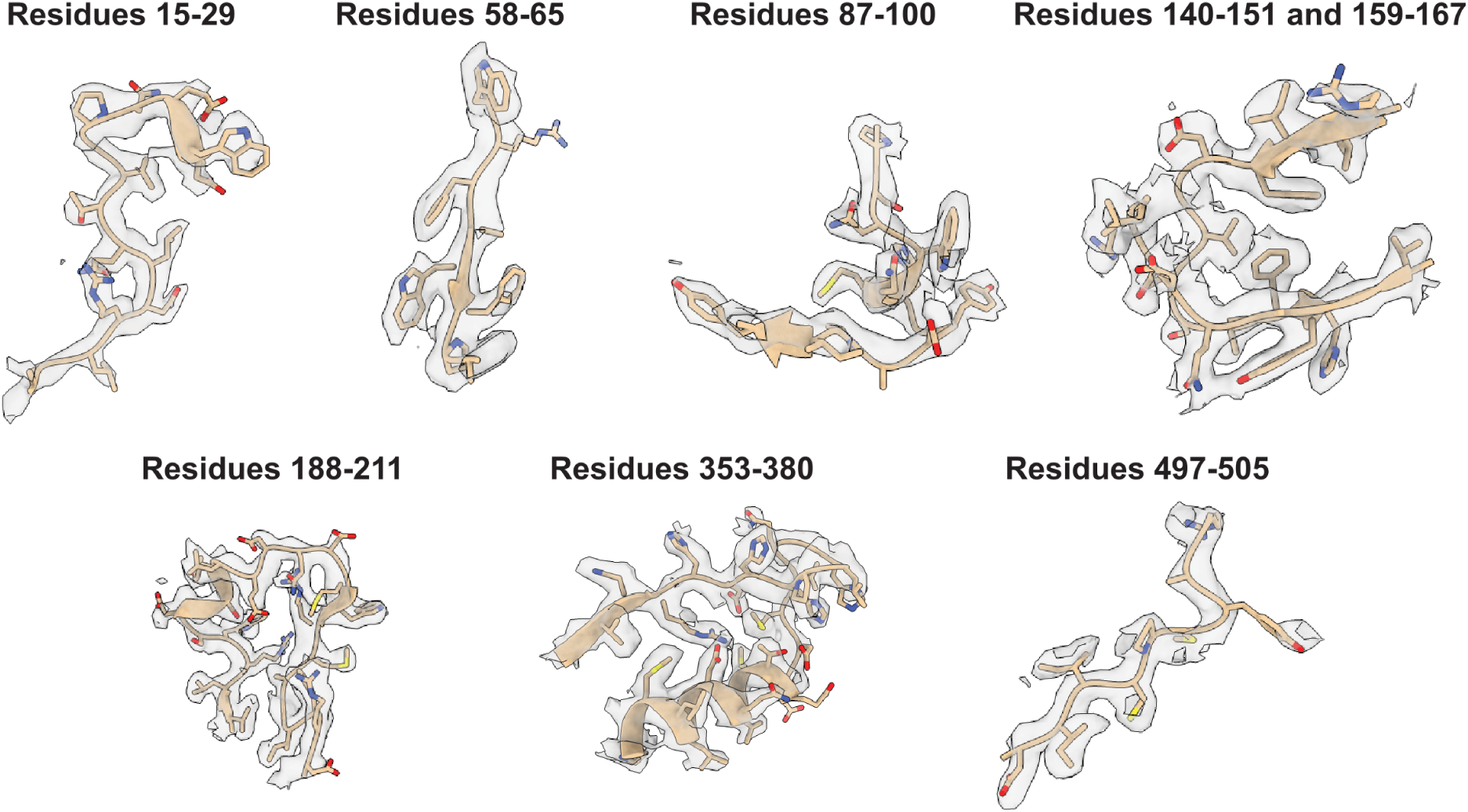
Fit of atomic model into native ESIBD map. Gallery of segments of the manually built model are shown within the cryo-EM density, showing resolution of side chains within the map.

**Figure S8:**
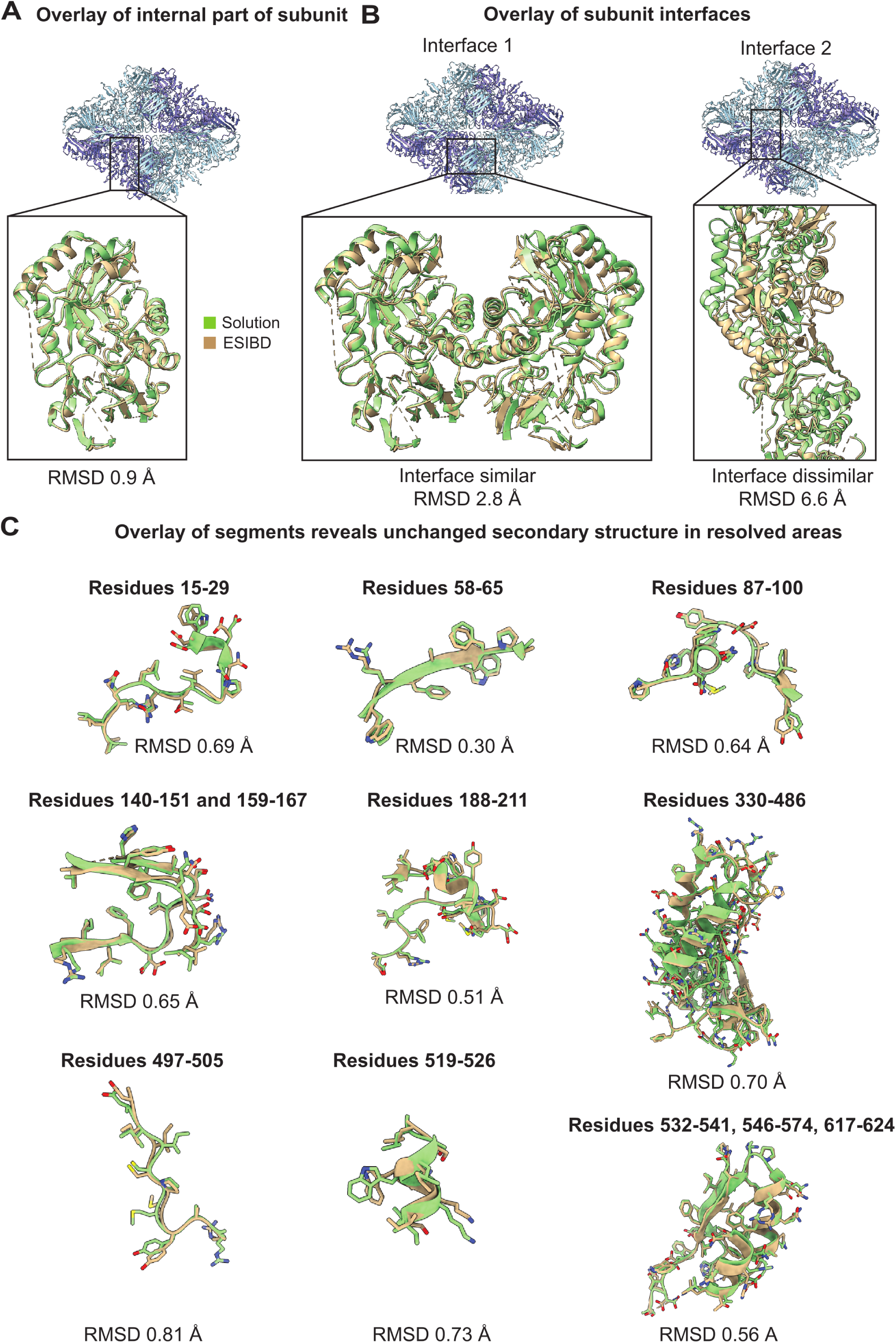
(Previous page) Comparison of built segments of native ESIBD *β*-galactosidase (tan) versus solution structure (green). **A** Comparison of a monomeric unit core region, aligned using the *matchmaker* function in ChimeraX. RMSD = 0.9 Å. **B** Left: Changes in the first type of interface between subunits in *β*-galactosidase. RMSD = 2.8 Å. Right: Changes in the second type of interface between subunits in *β*- galactosidase. RMSD = 6.6 Å. **C** Agreement between segments within one monomeric subunit, demonstrating conserved secondary structure. Segments were aligned and root mean square deviation was calculated using the *matchmaker* function in ChimeraX.

### 2.8 Computational Details

MD simulations were conducted with the GROMACS suite ^64^ (Version 2021.6), with the OPLS/AA forcefield and TIP4P water. The starting structure was prepared from the 6CVM PDB entry. For the dehydration simulations, the protein was prepared such that surface residues were protonated while buried residues were set to a protonation state at neutral pH. To this end, ionizable surface residues were identified using PropKa, ^65^ ordered according to their pKa values, and protonated until an overall charge state of +44 e (according to the average experimental charge) was reached. The protein was solvated with a 2 nm hydration shell, approx. 19500 water molecules, and centered in a cubic periodic box with an edge length of 50 nm. After steepest energy minimization the system underwent 63 simulation windows of 500 ps. Before each simulation window, all molecules and ions above a threshold distance of 2 nm from the molecule were removed, and velocities were re-initialized at 370 K. The leap-frog integrator with a timestep of 1 fs was used. Temperature was maintained constant at 370 K using the Nośe-Hoover thermostat with separate temperature coupling of protein and non-protein groups with a coupling constant of 0.5 ps. Bond lengths to H-atoms were constrained using the LINCS algorithm. Long-range Coulomb interactions were calculated using the particle mesh Ewald (PME) method, thus applying a neutralizing background charge.

For all simulations in bulk water, the protein was centered in a dodecahedral box with a 1 nm distance from the box edge. The system was solvated with TIP4P water and neu-tralized using 44 Cl-ions. After energy minimization with the steepest descent method, the system was equilibrated at 300 K for 100 ps using the velocity rescale thermostat with a time constant of 0.1 ps. Subsequently, the system was pressure equilibrated with the Parrinello-Rahman isotropic barostat with a time constant of 2 ps. Production runs at NPT conditions were run for 200 ns with a time step of 2 fs.

Representative structures were identified using the GROMACS clustering method gromos with an 0.15 nm RMSD cutoff. The centroids of clustered structures in the last 50 ns (solvation and rehydration) and 10 ns (dehydration), were chosen as the representative structures shown in Fig. 4. Furthermore, the number of H-bonds over the trajectory were calculated using GROMACS and the radius of gyration (ROG) was calculated with the Python package MDTraj (Version 1.9.7).

**Table S2:** Deviation between MD and experimental structures. The table provides C*α* RMSD values between structures shown in Fig. 4 in the main text, as obtained using the align command in ChimeraX. The deviations are in line with the changes in radius of gyration and number of hydrogen bonds shown in Fig. 4. However, the global RMSD does not capture the local similarities and deviations shown in Fig. S7 and Fig. S8.

| Compared structures | RMSD (Å) |
| --- | --- |
| Solvated MD vs. 6CVM (PDB) | 3.25 |
| Dehydrated MD vs 6CVM (relaxed to ESIBD map) | 5.16 |
| Rehydrated MD vs. 6CVM (PDB) | 4.62 |
| Solvated MD vs. Dehydrated MD | 7.18 |
| Solvated MD vs. Rehydrated MD | 5.31 |
| Dehydrated MD vs. 6CVM (PDB) | 5.78 |
| 6CVM (PDB) vs. 6CVM (relaxed to ESIBD map) | 7.55 |

**Figure S9:**
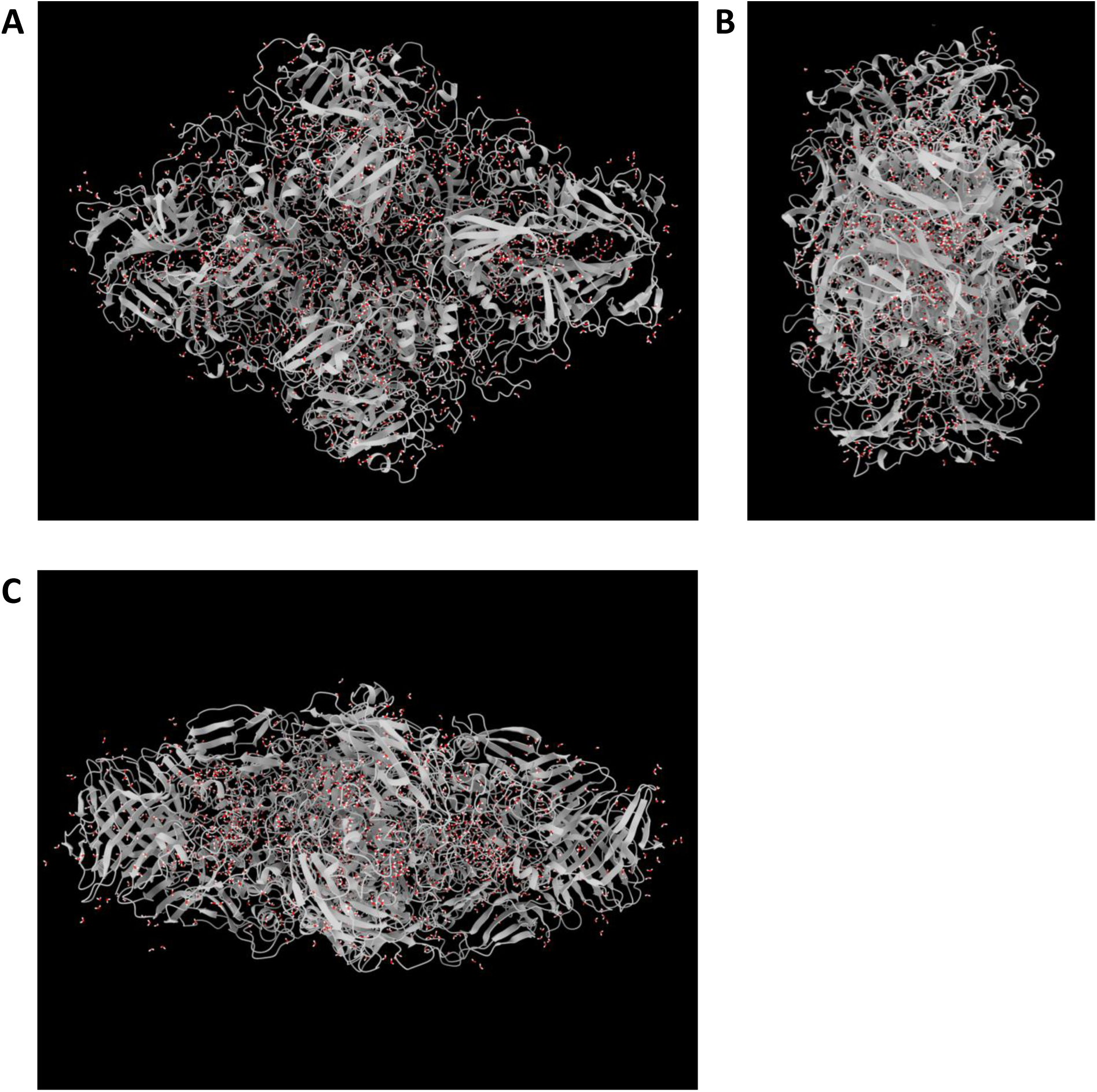
Position of water molecules in dehydrated MD structure. Most remaining 1,400 water molecules are found inside the protein structure. We have previously estimated the number of water molecules remaining in the experiment to less than 200.^36^ These 1,400 remaining water molecules are significantly less than the combined 4,194 structural water molecules found in the solvated structure (PDB 6CVM) and 4,200 molecules comprising a 3 Å solvation shell that could preserve the solvated structure.^24^ The different degrees of solvation in the dehydrated MD and native ESIBD structures can account for some of the structural differences between them. However, they both clearly represent significantly dehydrated states with respect to the native solvated structure.

